## Supplementary Figures for "Flexible Simultaneous Mesoscale Two-Photon Imaging Of Neural Activity at High Speeds"

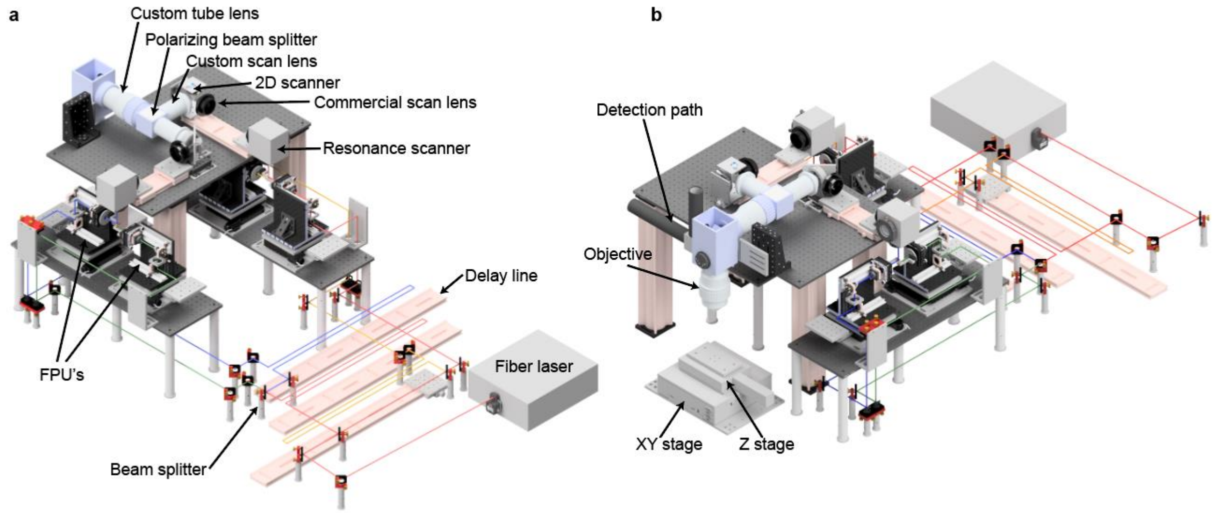


**Supplementary Figure 1. Mechanical design of the Quadroscope system. a)** View from the rear of the system. **b)** View from the front of the system.

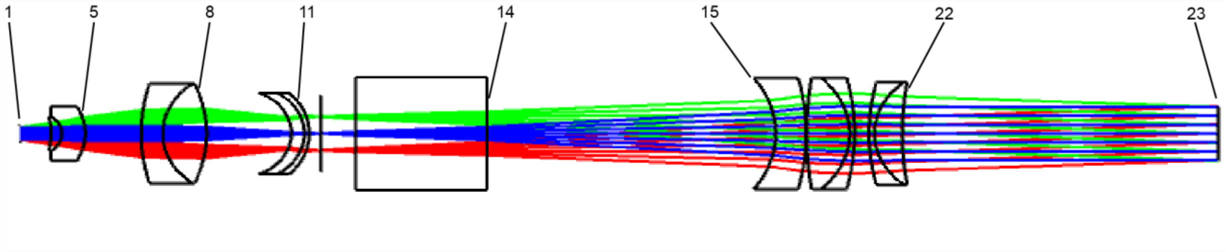


| Surface # | Description | Radius (mm) | Thickness (mm) | Material | Semi-Diameter (mm) |
|---|---|---|---|---|---|
| 1 | 2D scan mirror | Infinity | 15.499 | | 5.000 |
| 2 | | Infinity | 1.937 | | 10.743 |
| 3 | | -60.031 | 6.248 | S-PHM53 | 11.600 |
| 4 | | -13.894 | 13.411 | S-TIH53 | 11.600 |
| 5 | | -37.548 | 32.100 | | 18.000 |
| 6 | | 116.710 | 12.343 | S-TIH53 | 33.000 |
| 7 | | 40.204 | 25.041 | S-LAH53 | 33.000 |
| 8 | | -83.493 | 48.547 | | 33.000 |
| 9 | | -28.862 | 7.390 | S-TIH53 | 27.000 |
| 10 | | -31.235 | 3.000 | N-BK7 | 27.000 |
| 11 | | -37.279 | 6.600 | | 27.000 |
| 12 | | Infinity | 20.000 | | 25.702 |
| 13 | PBS | Infinity | 75.000 | N-BK7 | 37.500 |
| 14 | | Infinity | 165.233 | | 37.500 |
| 15 | | -60.635 | 17.500 | N-BK7 | 37.000 |
| 16 | | -145.360 | 0.250 | | 37.000 |
| 17 | | 250.339 | 24.511 | S-PHM52 | 37.000 |
| 18 | | -49.830 | 3.000 | LF5 | 37.000 |
| 19 | | -146.999 | 8.125 | | 37.000 |
| 20 | | 138.454 | 3.175 | LF5 | 34.000 |
| 21 | | 42.357 | 15.926 | S-FPL51 | 34.000 |
| 22 | | 207.680 | 181.098 | | 34.000 |
| 23 | Objective | Infinity | 0.000 | | 19.106 |

**Supplementary Figure 2. Lens specifications for custom scan lens and tube lens.**

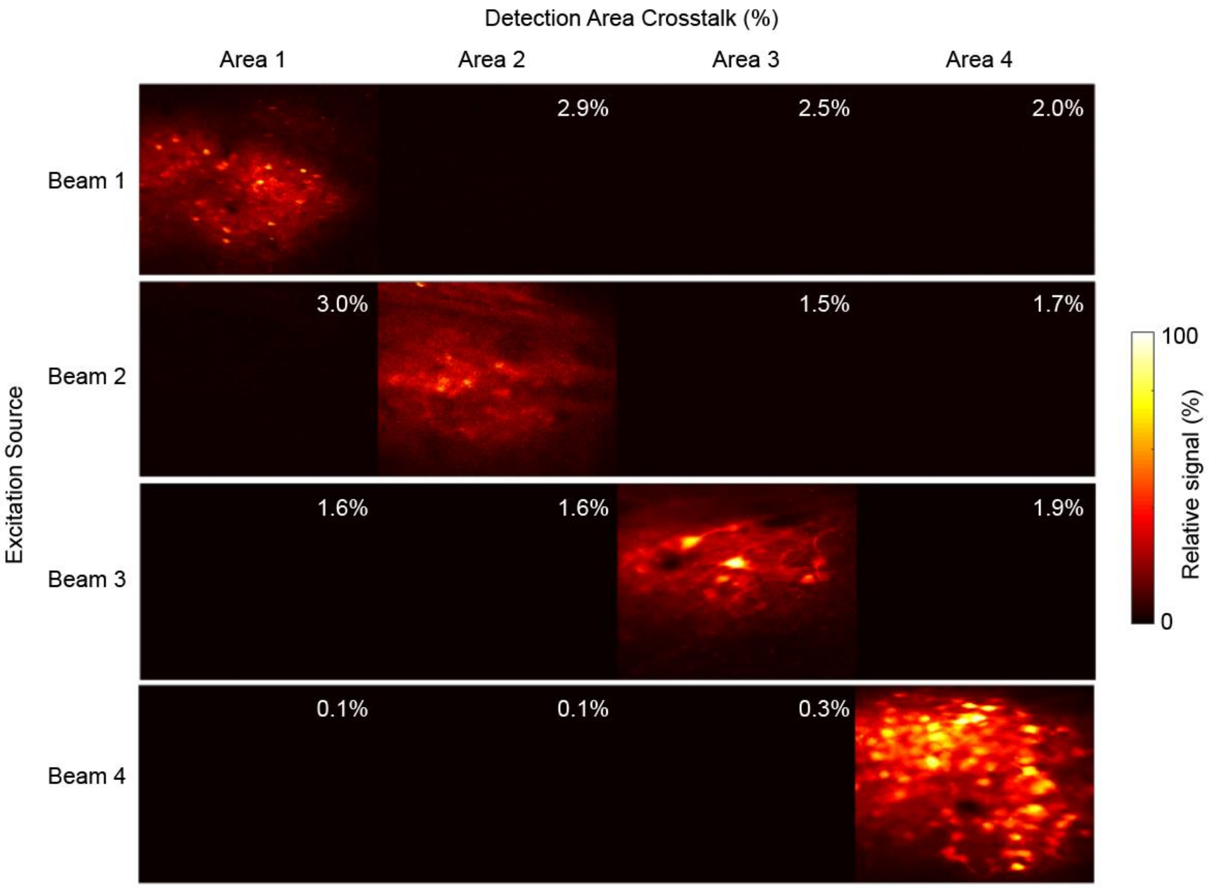


**Supplementary Figure 3. Crosstalk from Spatiotemporal multiplexing/de-multiplexing.** *In vivo* images of jYCaMP1s expression acquired sequentially from each excitation beam. De-multiplexed signals were assigned to each imaging area. Crosstalk for each area was calculated as the amount of detected signal relative to the main imaging area.

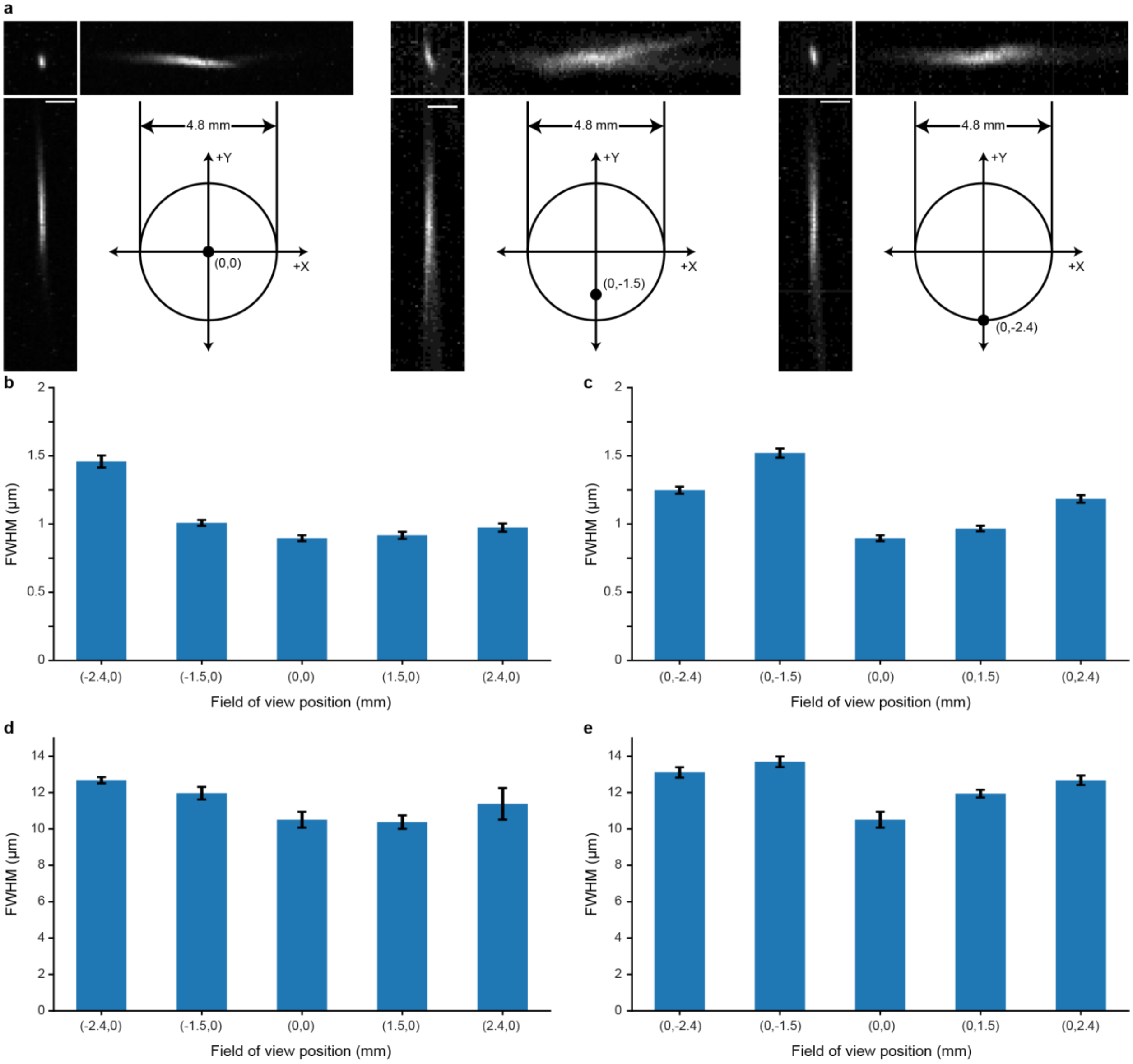


**Supplementary Figure 4. Point spread function measurements across the field of view. a)** Example images of 0.5 µm fluorescent beads from the center of the FOV, 1.5 mm off axis, and 2.4 mm off axis. Scale bars: 5 µm. **b,c)** Lateral full-width at half maximum (FWHM) measurements of beads along the full X axis [b] and the full Y axis [c] of the FOV. Bars represent the mean and error bars the s.e.m. A total of 8 beads were measured at each imaging location. **d,e)** Axial FWHM measurements of beads along the full X axis [d] and the full Y axis [e] of the FOV. The axial FWHM was measured in both the XZ and YZ planes to account for asymmetrical aberrations. Bars represent the mean and error bars the s.e.m.

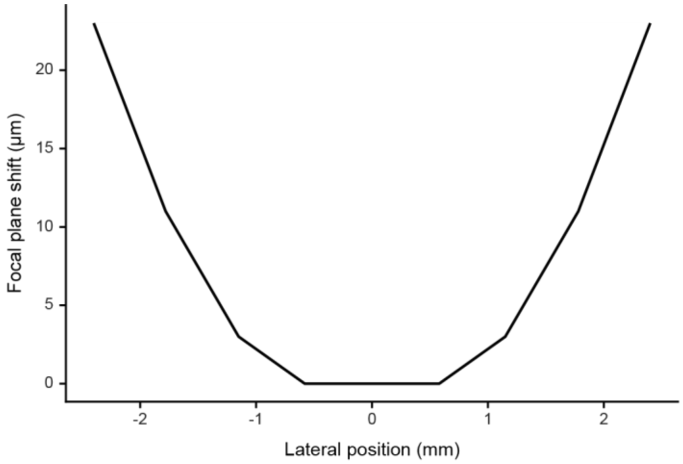


**Supplementary Figure 5. Field curvature.** Change in focal plane depth across the full FOV of the system as simulated in ZEMAX.

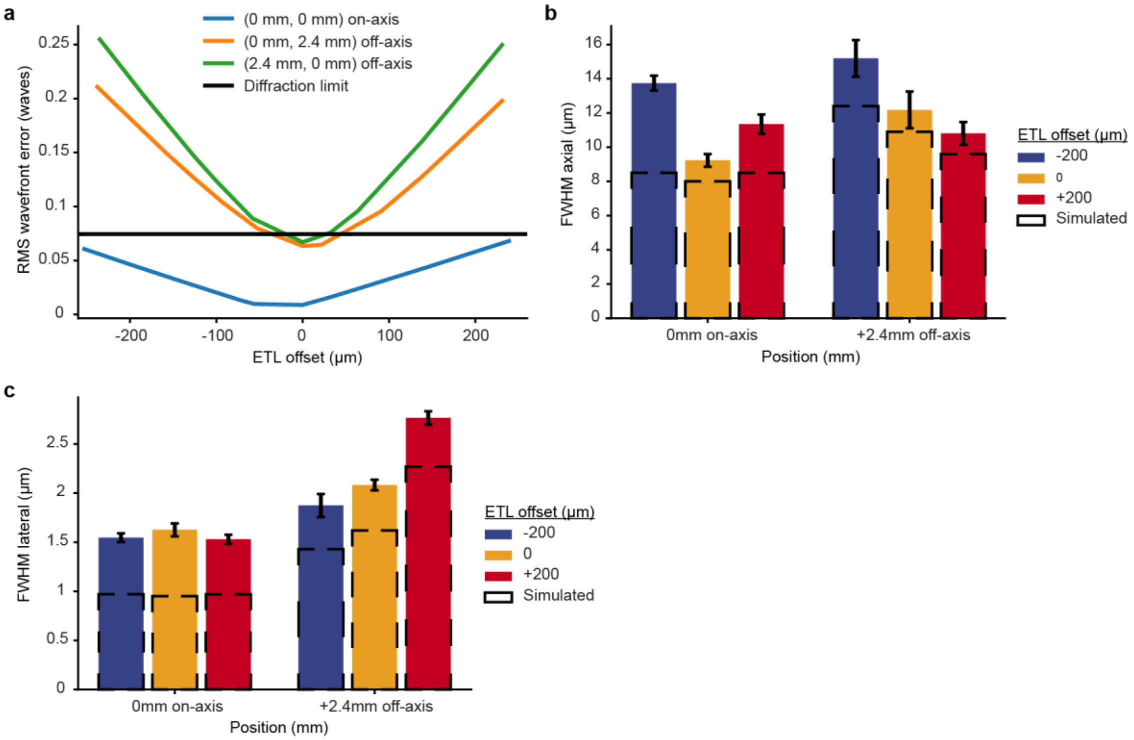


**Supplementary Figure 6. Performance impact of adjusting the ETL. a)** RMS wavefront error as a function of the focal plane change due to focusing of the electrically tunable lens (ETL). Performance is compared between the on axis condition and the 2.4mm off axis position in each the X and Y axis. **b and c)** Measured and simulated axial (**b**) and lateral (**c**) full width at half maximum PSF values. Measured values are from 0.5 µm fluorescent beads while simulated values come from the simulated Huygens PSF from ZEMAX. A total of 6-8 beads were collected for each imaging position. The bars represent the mean and error bars the s.e.m.

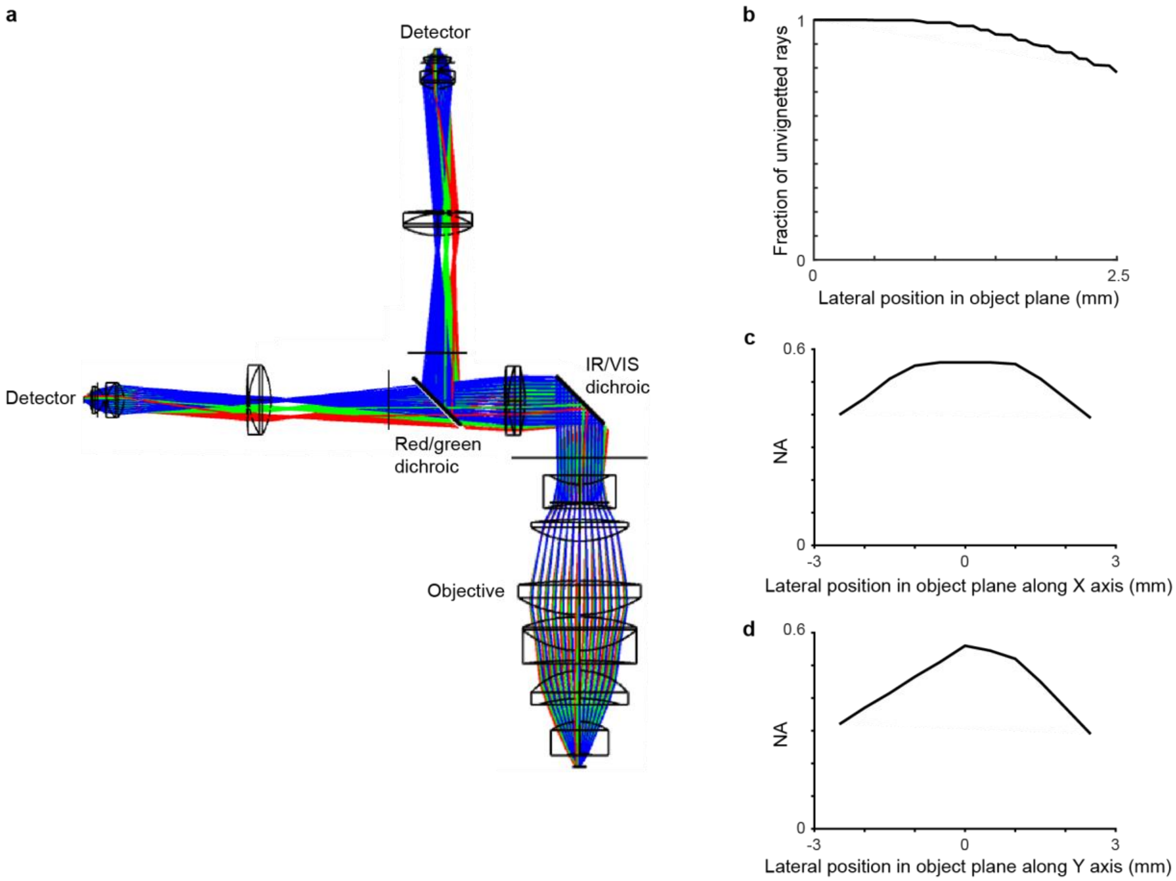


**Supplementary Figure 7. Optical design and analysis of the detection system. a**) Optical ray trace of the detection system. **b)** Plot of the fraction of modeled rays detected by the detector across the field of view. An efficiency of >78% was achieved across a 5 mm diameter FOV. **c and d**) Maximum numerical aperture at which all light reaches the detector as a function of position in the FOV along the x axis (**c**) and y axis (**d**).

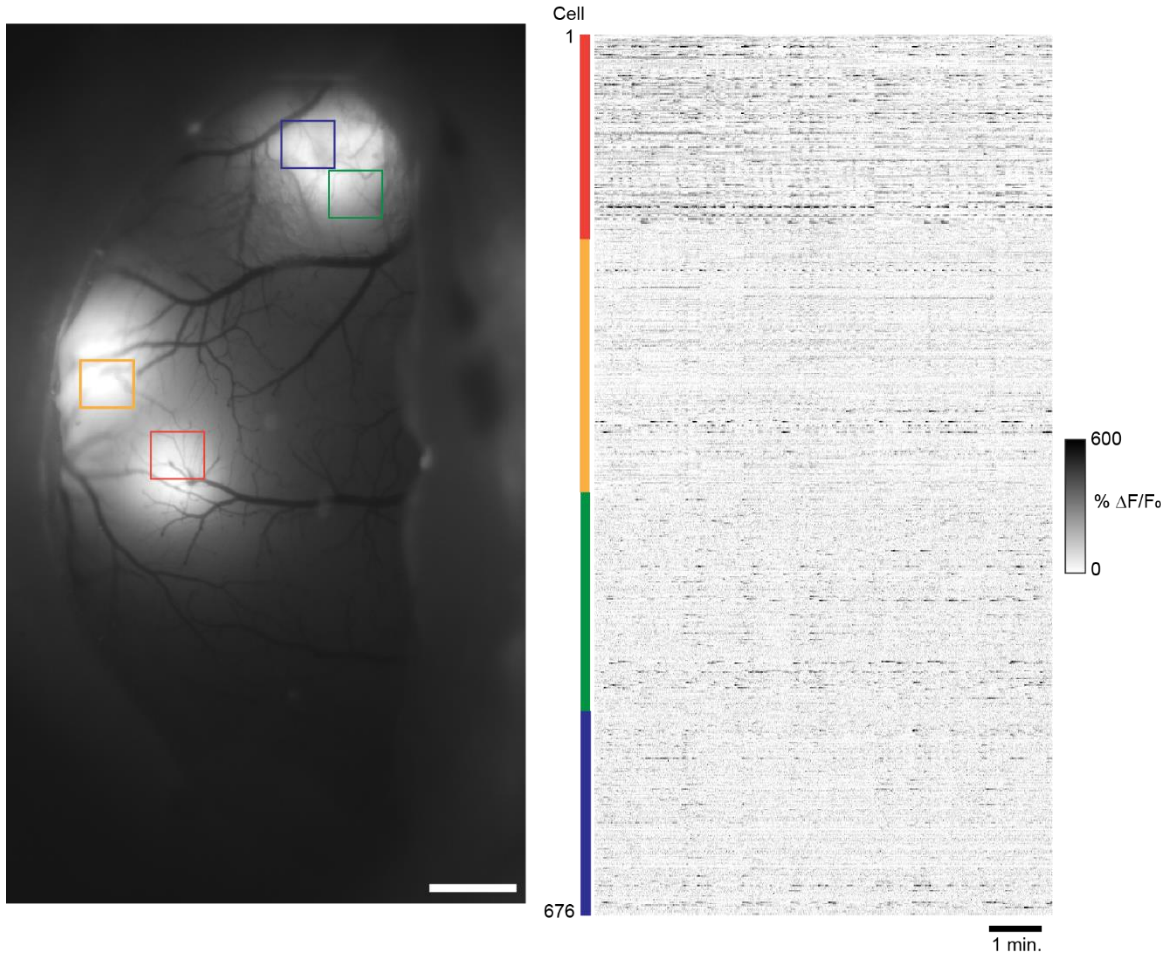


**Supplementary Figure 8. Quad area two-photon calcium imaging of GCaMP7f in sensorimotor cortex in the awake mouse using a 920nm laser source.** Wide-field fluorescence image of mouse sensorimotor cortex showing GCaMP7f expression through an implanted crystal skull (left). **b)** Calcium transients in the awake animal from cortical neurons imaged across the four areas, simultaneously acquired at 30 Hz (right). Scale bars: 1 mm.

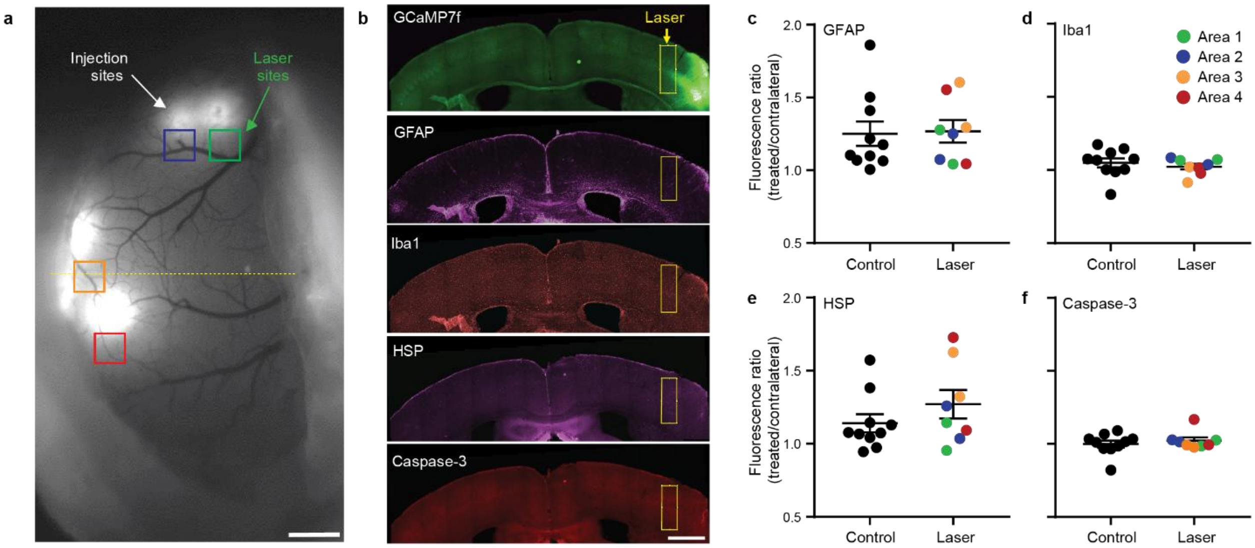


**Supplementary Figure 9. Sustained Quadroscope imaging does not induce photodamage. a)** Positioning of four imaging areas continuously scanned for 30 minutes at 75mW per beam .**b)** Examples of immunolabeled coronal sections on one imaging area (dotted line in [a]) with the area of laser exposure highlighted. **c-f)** No significant differences in relative fluorescence. between 8 laser-exposure areas (4 x 2 animals) and 10 control areas. Bars are mean ± s.e.m. Data points in the laser condition are color coded based on the four locations/beams.  Scale bars: 1 mm.
